# Developmental and genetic regulation of the human cortex transcriptome in schizophrenia

**DOI:** 10.1101/124321

**Authors:** Andrew E Jaffe, Richard E Straub, Joo Heon Shin, Ran Tao, Yuan Gao, Leonardo Collado Torres, Tony Kam-Thong, Hualin S Xi, Jie Quan, Qiang Chen, Carlo Colantuoni, William S Ulrich, Brady J. Maher, Amy Deep-Soboslay, The BrainSeq Consortium, Alan Cross, Nicholas J. Brandon, Jeffrey T Leek, Thomas M. Hyde, Joel E. Kleinman, Daniel R Weinberger

## Abstract

GWAS have identified 108 loci that confer risk for schizophrenia, but risk mechanisms for individual loci are largely unknown. Using developmental, genetic, and illness-based RNA sequencing expression analysis, we characterized the human brain transcriptome around these loci and found enrichment for developmentally regulated genes with novel examples of shifting isoform usage across pre- and post-natal life. We found widespread expression quantitative trait loci (eQTLs), including many with transcript specificity and previously unannotated sequence that were independently replicated. We leveraged this eQTL database to show that 48.1% of risk variants for schizophrenia associated with nearby expression. Within patients and controls, we implemented a novel algorithm for RNA quality adjustment, and identified 237 genes significantly associated with diagnosis that replicated in an independent case-control dataset. These genes implicated synaptic processes and were strongly regulated in early development (p < 10^-20^). These data offer new targets for modeling schizophrenia risk in cellular systems.

## Introduction

Schizophrenia (SCZD) is a prevalent neuropsychiatric disorder with a combination of genetic and environmental risk factors. Research over the last several decades has suggested that SZCD is a neurodevelopmental disorder arising through altered connectivity and plasticity in relevant neural circuits. However, discovering the causative mechanisms of these putatively developmental deficits has been very challenging^1^. The most consistent evidence of etiologic mechanisms related to SCZD has come from a recent genome-wide association study (GWAS) in which over a hundred independent single nucleotide polymorphisms (SNPs) were identified having a significant allele frequency difference between patients with schizophrenia and unaffected controls^2^. While these findings have identified regions in the genome harboring genetic risk variants, almost all of the associated SNPs are non-coding, located in intronic or intergenic sequence, and hypothesized to have some role in regulating expression^3^. However, the exact gene(s) and transcript(s) potentially regulated by risk-associated genetic variation are uncertain, as most of these genomic regions contain multiple genes. In principle, the effects of non-coding genetic variation, by whatever mechanisms (e.g. promoter, enhancer, splicing, noncoding RNA, epigenetics, etc), should be observed in the transcriptome. Therefore, to better understand how these regions of genetic risk and their underlying genotypes may confer risk of schizophrenia and to better characterize the molecular biology of the disease state, we sequenced the polyA+ transcriptomes from the prefrontal cortex of 495 individuals with ages across the lifespan, ranging from the second trimester of fetal life to 85 years of age (see Table S1), including 175 patients with schizophrenia (see Figure S1).

Here we identify novel expression associations with genetic risk and with illness state and explore developmentally regulated features, including a subset of genes with previously uncharacterized isoform shifts in expression patterns across the fetal-postnatal developmental transition. We further identify many more expression quantitative loci (eQTLs) in schizophrenia risk regions than previously observed by surveying the full spectrum of associated expression features to generate potential molecular mechanisms underlying genetic risk. We also explore differential gene expression associated with the state of illness in a comparison of the postmortem brains of patients with schizophrenia with non-psychiatric controls. By incorporating a novel, experiment-based algorithm to account for RNA quality differences which have not been adequately controlled in earlier studies, we report a high degree of replication across independent case-control gene expression datasets^4,5^. By combining genetic risk at the population-level with eQTLs and case-control differences, we identify putative human frontal cortex mechanisms underlying risk for schizophrenia and replicable molecular features of the illness state.

## Results

We performed deep polyA+ RNA-sequencing of 495 individuals, ranging in age from the second trimester of fetal life to 85 years old (see Table S1), including 175 patients with schizophrenia (see Figure S1). We quantified expression across multiple transcript features, including: annotated 1) genes and 2) exons, 3) annotation-guided transcripts^4^ as well as alignment-based 4) exon-exon splice junctions^5^ and 5) expressed regions (ERs)^6^. These last two expression features were selected to reduce reliance on the potentially incomplete annotation of the brain transcriptome^7^ (Results S1). We find a large number of moderately expressed and previously unannotated splice junctions that tag potential transcripts with alternative exonic boundaries or exon skipping (Figure S2), 95% of which are also found in other large RNA-seq datasets, including a subset that were brain-specific (Table S2). Similarly, we find that only 56.1% of ERs were annotated to strictly exonic sequence – many ERs annotated to strictly intronic (22.3%) or intergenic (8.5%) sequence, or were transcribed beyond existing annotation (e.g. extended UTRs, extended exonic sequence).

### Developmental regulation of transcription and shifting isoform usage

Characterizing expression changes in unaffected individuals, particularly across brain development beginning with prenatal life, has previously offered disease-relevant insights into particular genomic loci ^8-12^. Specifically, we and others ^7,13,14^ have shown that genomic risk loci associated with neurodevelopmental disorders including schizophrenia are enriched for transcript features showing differential expression between fetal and postnatal brains. Here too, among the 320 control samples, the strongest component of expression change corresponded to large expression changes in the contrast of pre-natal and early postnatal life, in line with previous data ^7^ (Figure 1A). We further defined a developmental regulation statistic for each expressed feature using a generalized additive model (see Methods) and found widespread developmental regulation of these expressed features (Results S2, Table S3, Figure S3), including previously unannotated sequence (Table S4). Motivated in part by previous reports of preferential fetal isoform use among schizophrenia candidate risk genes^10,11^ (e.g predominant fetal versus predominant postnatal isoforms), we next formally identified the subset of genes showing alternative isoform expression patterns across fetal and postnatal life using those exons, junctions, transcripts, and ERs that meet the statistical criteria for developmental regulation (i.e. those genes with at least one developmentally changing feature, see Methods). We highlight a representative gene with isoform shifts in Figure S4 involving *CRTC2*, a transcription co-activator. There were 6672 Ensembl genes (23.7% of the set of developmentally regulated genes) with both positive and negative expression features having genome-wide significant correlations with age (each with p_bonf_<0.05, Figure 1B, Table S5, Figure S5). In other words, these represent alternate transcript isoforms of the same gene that show opposite patterns of expression across the prenatal-postnatal transition. In principle, this interaction would obscure developmental expression variation measured at the gene level.

**Figure 1:**
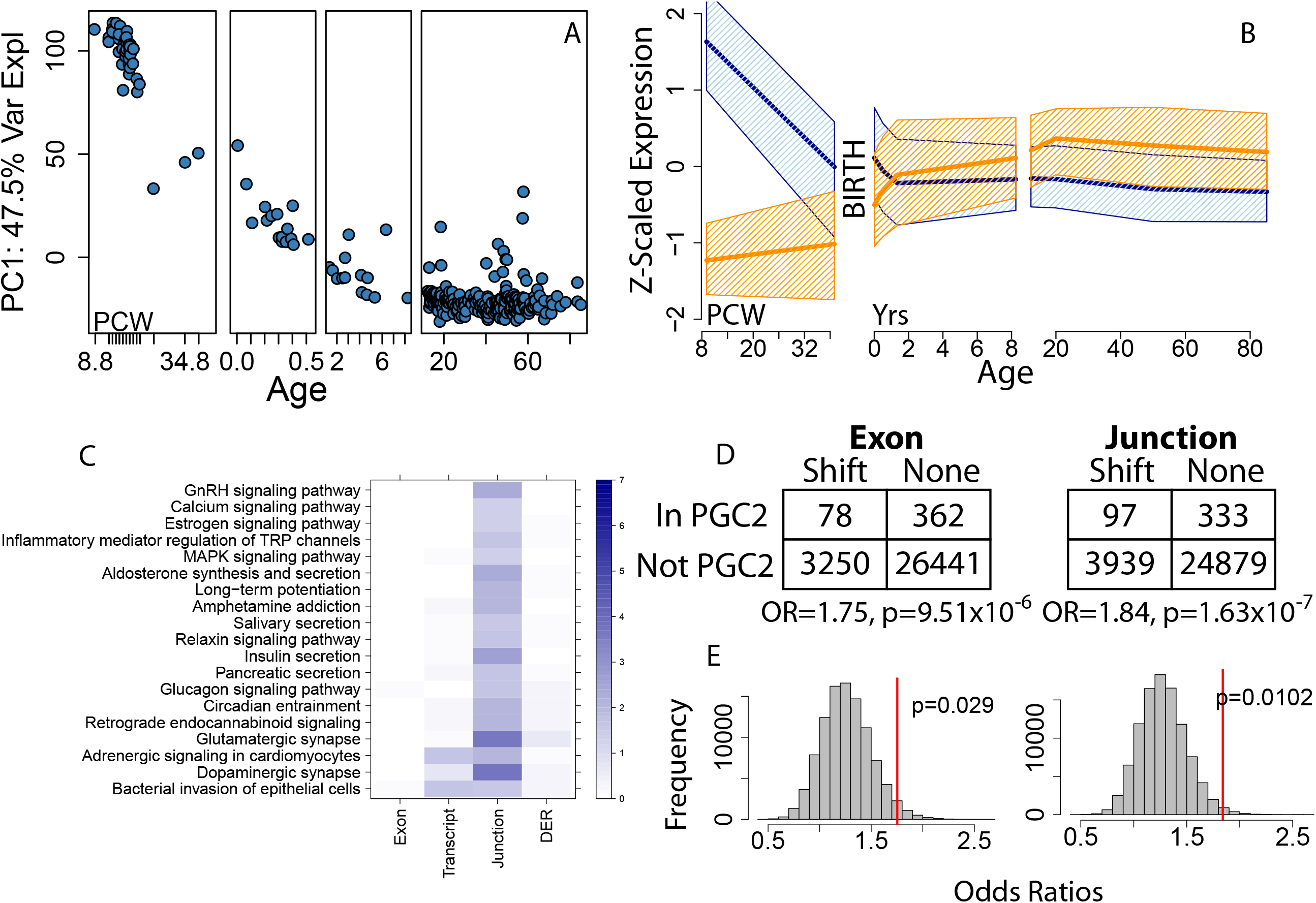
Developmental regulation of expression. (A) Principal component #1 of the gene-level expression data versus age; PCW: post-conception weeks, remaining ages are in years. (B) Expression features fall into two main development regulation signatures, increasing in expression from fetal to postnatal life (orange) or decreasing from fetal to postnatal life (blue). Y-axis is Z-scaled expression (to standard normal), dark lines represent median expression levels, and confidence bands represent 25^th^-75^th^ percentiles of expression levels for each class of features. (C) KEGG pathways enriched for genes with isoform shifts, stratified by which feature type identified the gene as having a switch. Coloring/scaling represents -log10(FDR) for gene set enrichment. Analogous data for GO gene sets (biological processes, BP, and molecular function, MF) are available in Table S6. DER: differentially expressed region. Enrichment analyses for isoform shift genes among PGC2 schizophrenia GWAS risk loci with exon and junction counts using both (D) parametric p-values) and (E) permutation-based p-values. OR: odds ratio.

We performed gene set analyses of those genes with shifting isoform usage compared to the larger set of genes with at least one developmentally regulated feature but without shifting isoform usage to identify more specific biological functions of this unique form of developmental regulation (Table S6). The set of developmentally shifting isoforms was relatively enriched for localization, catalytic activity, signaling-related processes, including synaptic transmission and cell communication, and neuronal development, among many others. Interestingly, genes identified with shifting isoforms across development based exclusively on junction counts were enriched for both dopaminergic (FDR=1.67x10^−4^) and glutamatergic (FDR=2.04x10^−4^) synapse KEGG pathways (Figure 1C), the two neurotransmitter systems most prominently implicated in schizophrenia pathogenesis and treatment.

### Schizophrenia risk is associated with novel shifting isoform usage across brain development

Based on the KEGG analysis, we hypothesized that the genes with developmentally regulated isoform shifts may relate to risk for schizophrenia. Indeed, genes within the SCZD GWAS risk loci were more likely to harbor these novel isoform shifts occurring in the fetal-postnatal developmental transition compared with the rest of the expressed transcriptome (Figure 1D). For example, genes with developmental isoform shifts identified by exon, junction and expressed region counts were 75% (p=9.51x10^−6^), 84% (p=1.63x10^−7^) and 71% (p=2.0x10^−4^) more likely to lie within the PGC2 risk regions (with permutation-based p=0.02, p=0.01, and p=0.03 respectively, see Methods) than developmentally regulated genes without isoforms shifts (Table S7). These results further underscore the role of changes in the regulation of transcription and splicing in the early brain developmental components of schizophrenia risk.

### Large-scale genetic regulation of transcript-specific and previously unannotated sequences

In order to elucidate the RNA features associated with schizophrenia risk variants themselves, rather than positional LD regions, we first performed a genome-wide *cis* (<500kb) expression quantitative trait loci (eQTL) analysis within the 412 post-adolescent subjects (see Methods) across the five convergent transcript features (Table 1). We hypothesized that, in general, analyzing transcript features like exons and junctions would increase statistical power for eQTL discovery if genetic variation regulated the expression levels of specific mRNA transcripts. At the gene-level, which collapses data from all transcripts into a single measure, which is the most common feature summarization for eQTL discovery, the vast majority of expressed genes were associated with the expression of at least one nearby genetic variant. There were eQTLs to 6748 Ensembl Gene IDs (of which 4955 genes had HGC symbols) at stringent Bonferroni-adjusted significance (p < 8.41x10^−9^, see Methods), and eQTLs to 18,416 Ensembl Gene IDs at more liberal FDR < 1% significance (p < 1.84x10^−4^). However, we found a larger number of genes with eQTLs using exon-level analysis – 48,031 exons mapping to 8386 Ensembl IDs - at Bonferroni significance (“eExons”, p < 7.64x10^−10^). Exon-level analysis showed widespread transcript-specificity of eQTL associations. Almost all eExons mapped to genes with more than one annotated transcript (N=45,239, 94.2%), and the majority of these showed eQTL associations to exons belonging to a single transcript isoform (N=30,283, 66.9%). This transcript-specificity was also evident in the eQTL effect sizes, as the median additive effect size was approximately two-fold higher for exon- than gene-level analysis (15.6% versus 7.0% expression change per allele copy). Interestingly, while transcript-specific by nature, we actually found the fewest eQTLs to assembled-and-quantified transcripts (3,263 eTxns at p < 1.73x10^−9^), in line with previous reports highlighting the difficulties in merging assemblies across replicates^15^. Lastly, there were an additional 3,022 eGenes identified with exon-level analysis compared to the 5364 eGenes identified with both summarization levels. These results demonstrate extensive transcript specificity of many eQTL signals that are missed by gene-level analyses.

**Table 1:** eQTL summary statistics at FDR and Bonferroni significance thresholds across five feature summarizations. “logFC” is the log2 fold change in expression per minor allele copy and “% Unann” is the percent of features that were not strictly annotated.

|  | Type | eQTLs | # SNPs | # Features | p-cutoff | Ensembl Genes | Symbol Genes | log2FC | % Unann |
| --- | --- | --- | --- | --- | --- | --- | --- | --- | --- |
| FDR < 1% | Gene | 1815172 | 1055186 | 18416 | 1.84E-04 | 18416 | 12874 | 0.061 | NA |
|  | Exon | 13255860 | 1390362 | 157923 | 1.00E-04 | 20696 | 15697 | 0.13 | NA |
|  | Transcript | 1465179 | 616346 | 26870 | 3.07E-05 | 11272 | 11219 | 0.094 | 50.7% |
|  | Junction | 4813472 | 1092615 | 67358 | 6.39E-05 | 14792 | 13204 | 0.33 | 21.3% |
|  | ER | 8115891 | 1367619 | 94200 | 1.25E-04 | 16379 | 12914 | 0.22 | 47.4% |
| Bonf < 5% | Gene | 648597 | 431704 | 6748 | 8.41E-09 | 6748 | 4955 | 0.097 | NA |
|  | Exon | 4019197 | 529237 | 48031 | 7.64E-10 | 8386 | 6439 | 0.21 | NA |
|  | Transcript | 514563 | 236633 | 6349 | 1.73E-09 | 3263 | 3249 | 0.15 | 46.9% |
|  | Junction | 1557370 | 439920 | 18908 | 1.10E-09 | 5827 | 5205 | 0.55 | 21.6% |
|  | ER | 2575655 | 533978 | 27643 | 1.28E-09 | 6822 | 5643 | 0.37 | 53.9% |

We next explored the extent of eQTLs to previously unannotated transcribed sequence using junction- and expressed-region feature summarizations which do not rely on existing gene annotation for quantification. Among the 18908 junctions with eQTL signal at Bonferroni significance (“eJxns”, p<1.1x10^−9^), 21.6% (N=4089) were previously unannotated, including 1312 eJxns to exon-skipping splicing events and 2777 eJxns to shifted exonic boundaries (acceptor or donor splice sites). The eJxns also highlight a large degree of potential transcript specificity, both in the 4089 unannotated junctions as well as 3388 additional annotated eJxns that delineate individual transcript isoforms (when multiple isoforms are present for the gene). At the expressed region-level, among the 27,643 ERs with eQTL signal at Bonferroni significance (“eERs”, p<1.28x10^−9^), 14,890 were either fully or partially unannotated, with partial events including 4521 exon extensions into neighboring intronic sequence and 769 extended untranslated regions (UTRs) and fully unannotated events being strictly intronic (N=6,255) and intergenic (N=3,345) sequences. These two feature classes also had the largest eQTL effect sizes of the tested features, with 41.4% and 29.2% change in expression per allele copy for eJxns and eERs. Lastly, we found that 1,042 Ensembl genes had eQTLs exclusively to unannotated sequence with no corresponding eQTL signal to annotated features in the genes. Genetic regulation of previously unannotated sequence provides further evidence for biological relevance in the human brain.

Given the large degree of genetic regulation of transcript-specificity and unannotated sequences, we sought to assess the replication of the identified eQTLs (“LIBD”) in independent human brain RNA-seq data. We downloaded alignment-level data from the CommonMind Consortium (“CMC”) project, and quantified expression across the same five feature summarizations (in Table 1). Among those significant eQTL SNP-feature pairs that were well-imputed, polymorphic and expressed in the replication dataset (~84% of pairs, ~95% of eFeatures, see Methods, Figure S6), >94% had consistent directionality in the two datasets, between 75.7% (eTxns) and 81.5% (eJxns) were directionally consistent and marginally significant (at p < 0.01), and just over half (52.1%-57.0%) were directionally consistent and FDR-corrected significant (published set, p<10^−5^) in the DLPFC replication dataset. Meta-analysis between datasets demonstrated extensive significance and replication of the 9.3M SNP-feature Bonferroni-significant eQTL pairs including 97.6% at p < 1x10^−5^ and 82.0% at p <10^−9^. We further reprocessed and quantified GTEx v6 RNA-seq brain data (“GTEx”) from raw reads using the same pipeline, and assessed replication and regional specificity in these data using meta-analysis across 13 brains regions compared to frontal cortex alone. Here we found that many of the DLPFC-identified eQTLs showed strong concordant signal across all brain regions, suggesting an overall lack of regional specificity for the majority of our identified eQTLs (Figure S7). All significant eQTLs are searchable on our publicly available database: eqtl.brainseq.org/phase1/eqtl/ which provides visualizations and eQTL statistics across three independent datasets.

### Clinical enrichment of eQTL associations for schizophrenia and other traits

We sought to better determine the clinical relevance of our significant eQTLs particularly in the context of transcript feature-level and previously unannotated sequence associations. We cross-referenced our identified eQTLs with genome-wide association study (GWAS) risk variants. Here we used 3 significance levels to associate eQTLs with GWAS variants: a) more liberal FDR-significant eQTLs in the discovery dataset, b) these FDR-significant eQTLs with additional replication data support (meta-analysis p-values with CMC < 10^−8^), and c) Bonferroni-significante QTLs in the discovery dataset, e.g. Table 1, First we considered the proportion of common (MAF > 5%) and well-measured risk variants from the 128 index variants (N=106, see Methods) published in the latest PGC2 GWAS for schizophrenia ^2^ and their highly correlated proxies (see Methods). We identified FDR-significant eQTL associations to 51 risk SNP signals (of 106 tested, 48.1%, Table S8), a substantially higher proportion of risk variants classified as brain eQTLs than previously reported^16^ (Table 2). In total, there were 1,244 unique SNP-feature pairs that were genome-wide FDR-significant eQTLs (83 genes, 553 exons, 49 transcripts, 192 junctions and 367 ERs) mapping to 194 unique Ensembl Gene IDs (of which 162 have HUGO gene symbols). Among these 51 risk SNPs, 17 were eQTLs only to exons, junctions or expressed regions, and 7 were eQTLs to only unannotated transcribed sequence. There were 17 loci with annotated eQTLs to only a single gene and another 10 loci with eQTLs to two genes. More stringent meta-analysis significance (p<10-8) retained eQTL evidence for 37 variants including 17 to exons, junctions, and ERs, of which 6 were unannotated.

**Table 2:** eQTL summary metrics for GWAS variants from the latest schizophrenia GWAS and the more general genome-wide suggestive loci from the NHGRI GWAS catalog. “# SNPs Tested” were those that were observed or imputed with high quality and that were relatively common in our samples (MAF > 5%). “Unann” = unannotated, “Tx” = transcript

|  | SCZD GWAS |  |  | NHGRI GWAS Catalog |  |  |
| --- | --- | --- | --- | --- | --- | --- |
|  | FDR<1% | FDR+Meta | Bonf<5% | FDR<1% | FDR+Meta | Bonf<5% |
| # SNPs Tested | 106 | 106 | 106 | 23704 | 23704 | 23704 |
| # SNP eQTLs | 51 | 37 | 26 | 8988 | 5490 | 4255 |
| > # w/o Gene | 21 | 17 | 9 | 3763 | 2370 | 1891 |
| > # w/o Gene+Tx | 17 | 15 | 8 | 2982 | 1824 | 1445 |
| > # Unann | 47 | 28 | 17 | 5858 | 3470 | 2579 |
| > # Only unann | 7 | 6 | 3 | 995 | 671 | 589 |
| > # Single Tx | 11 | 10 | 5 | 1933 | 1156 | 976 |

We also assessed enrichment of 23704 GWAS risk SNPs from the NHGRI GWAS catalog present and common in our genetic data (of 44,738 available), and found eQTL evidence for 8988 variants (37.9%) at FDR < 0.01. These GWAS variants that were identified as eQTLs were from GWAS for the majority of all tested traits in the literature (68.1%, 1415 of 2078 present) across all sites in the body, suggesting that many of the identified eQTLs in brain are likely shared with other tissue sites as previously described ^17^. Of the 8988 GWAS eQTL variants, 2982 were eQTLs only to exons, junctions or expressed regions, of which 995 were only to unannotated sequence (Table 2). More stringent meta-analysis significance (p<10^−8^) retained eQTL evidence for 5490 variants including 1824 to exons, junctions, and ERs, of which 671 were unannotated. These results highlight the ability to identify more eQTL signal for clinical risk variants by casting a wider net of RNA-seq feature summarization, including previously unannotated transcribed sequences.

### Refining risk transcripts through conditional analyses

We further sought to filter the eQTL hits to schizophrenia GWAS regions using conditional analysis in order to identify perhaps the most immediate downstream features of genetic risk. For each of the 51 eQTL-positive GWAS variants noted above, we conditioned on the most significant eQTL feature for each variant and then performed eQTL reanalysis of all other features. We then retained those eQTL features that remained at least marginally significant (at p <0.05) and repeated the conditional analysis now based on the two most independently associated expression features. We iteratively performed these conditional analyses until no other features were conditionally significantly associated eQTLs. These analyses resulted in only 220 conditionally-independent SNP-feature eQTLs (35 genes, 66 exons, 8 transcripts, 50 junctions and 61 ERs) to the 51 schizophrenia GWAS variants (Table S8) which mapped to 131 unique Ensembl Gene IDs (of which 106 have HUGO gene symbols). Conditional analysis resulted in an additional locus with eQTLs to a single gene (totaling 18 loci) and an additional four loci with eQTLs to features in two genes (totaling 14 loci, Table 3). Interestingly, these conditional analyses further highlighted the potential importance of transcript-specific and previously-unannotated eQTLs, as more loci were associated only with exons, junctions and ERs (27 versus 17), more were strictly unannotated (11 versus 7), and more showed eQTL associations to a single transcript isoform (18 versus 11).

**Table 3:** GWAS-significant index variants and eQTL associations, for those GWAS loci associating with only one or two genes following conditional analysis.

| SZ GWAS Locus | SNP | Gene | SZ GWAS Locus | SNP | Gene |
| --- | --- | --- | --- | --- | --- |
| 1 | rs1233578 | Intergenic | 59 | rs10520163 | *CLCN3* |
| 1 | rs1233578 | *ZSCAN26* | 63 | rs9420 | Intergenic |
| 5 | rs4129585 | *TSNARE1* | 73 | rs3849046 | *CTNNA1* |
| 7 | rs10650434 | *MAD1L1* | 82 | rs6704641 | *SATB2* |
| 7 | rs10650434 | *FTSJ2* | 84 | rs1106568 | *GPM6A* |
| 11 | rs4702 | *FES* | 86 | rs10043984 | *FAM53C* |
| 11 | rs4702 | *AC068831.1* | 86 | rs10043984 | *NME5* |
| 12 | rs75968099 | *LRRFIP2* | 88 | rs7819570 | *AC090568.2* |
| 12 | rs75968099 | *AC011816.1* | 96 | rs8082590 | *ATPAF2* |
| 16 | rs13240464 | *LRRN3* | 96 | rs8082590 | *DRG2* |
| 16 | rs13240464 | *IMMP2L* | 98 | rs12325245 | *GOT2* |
| 17 | rs10791097 | *SNX19* | 98 | rs12325245 | *NDRG4* |
| 20 | rs7893279 | *NSUN6* | 103 | rs324017 | *STAT6* |
| 23 | rs6704768 | *C2orf82* | 106 | rs9841616 | *SOX2-OT* |
| 23 | rs6704768 | *GIGYF2* | 109 | rs149009306 | *DFNA5* |
| 24 | rs55661361 | *NRGN* | 114 | rs12421382 | *AP003049.1* |
| 30 | rs11682175 | *FANCL* | 114 | rs12421382 | Intergenic |
| 42 | rs7432375 | *AC117382.2* | 117 | rs75575209 | *FANCL* |
| 42 | rs7432375 | *PCCB* | 119 | rs14403 | *AKT3* |
| 47 | rs4523957 | *SRR* | 119 | rs14403 | *SDCCAG8* |
| 47 | rs4523957 | *TSR1* | 120 | rs6670165 | *BRINP2* |
| 52 | rs140505938 | Intergenic | 120 | rs6670165 | Intergenic |
| 57 | rs34269918 | *RERE* | 121 | rs7523273 | *CD46* |
| 57 | rs34269918 | *SNORA77* |  |  |  |

We highlighted several representative eQTLs in Figure 2 for different classes of associations. The top GWAS risk variant rs1233578 associated with strictly intergenic sequence downstream of *ZSCAN23* (Figure 2A,2B, p=2.7x10^−8^) with replication in both CMC (p=0.01) and GTEx (T=3.1), suggesting potential novel transcribed sequence linked to schizophrenia risk. We also found significant eQTL signal to specific 5’ junction and exon sequences of *CTNNA1* to rs3849046 (Figure 2C,2D; discovery p=6.2x10^−8^, CMC replication p=1.4x10^−8^). Another example of eQTL associations of partially annotated sequence was rs9841616 exclusively associating with the 3’ sequence of the most proximal short transcript isoform of *SOX2-OT* (Figure 2E,2F; discovery p=8.2x10^−12^, replication p=2.9x10^−8^). We also found novel eQTL associations to annotated exons in *CD46* (Figure 2G, p=9.2x10^−38^, replication p = 2.9x10^−14^), *SRR* (Figure 2H,p=2.0x10^−12^, replication p=4.7x10^−6^) and *GPM6A* (Figure 2I, p=2.8x10^−6^, replication p=0.02).

**Figure 2:**
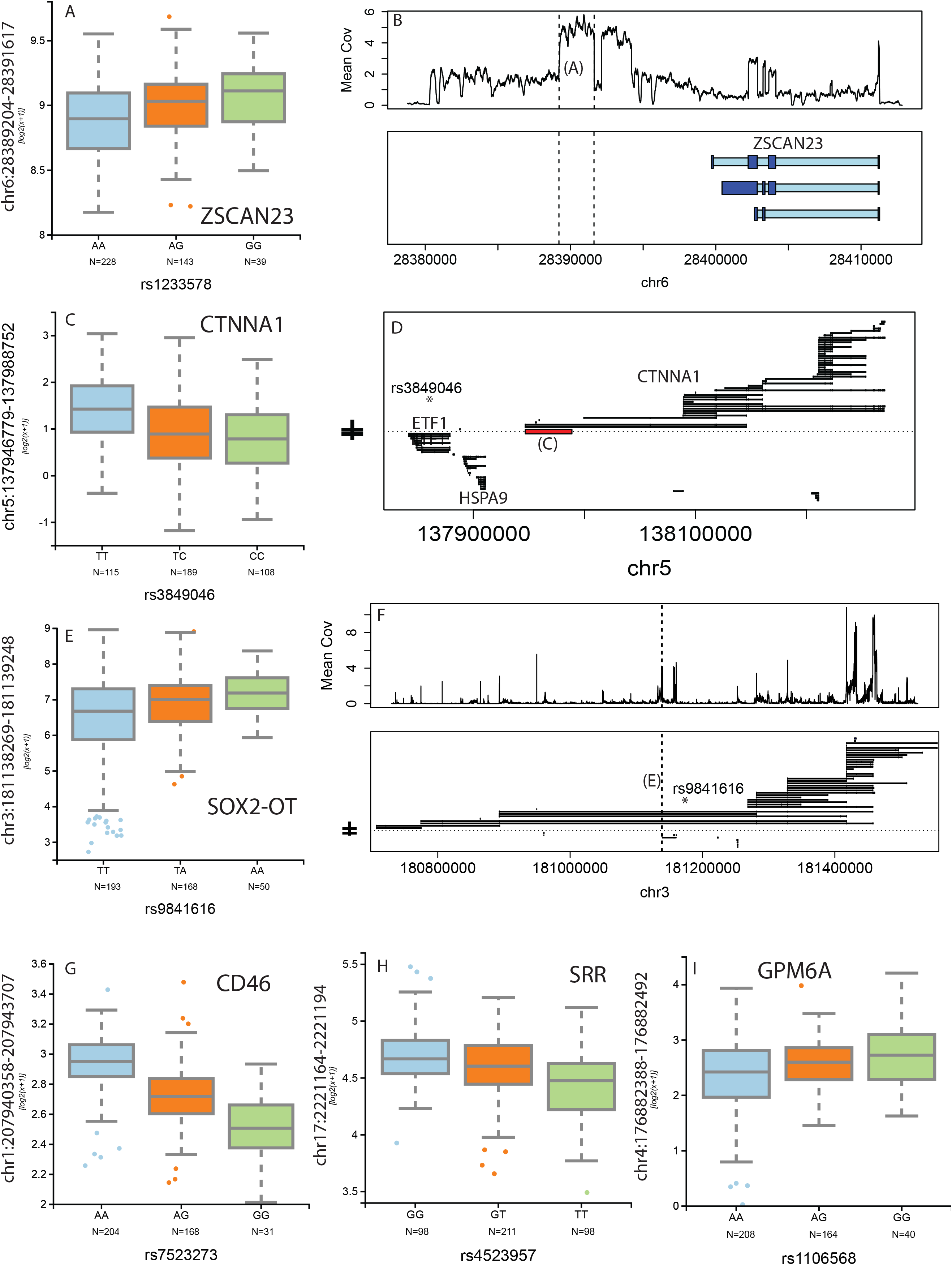
Clinical enrichment of schizophrenia risk using representative eQTLs. (A) Association between rs1233578 and intergenic sequence downstream (B) of ZSCAN23. (B) Association between rs3849046 and a splice junction (C) of a particular longer isoform (D) of *CNNTA1*. (E) Association between rs9841616 and very proximal extended UTR (F) of SOX2-OT Associations between risk SNPs and annotated sequences are shown for (G) CD46, (H) SRR and (I) GPM6A. In panels B, D, and F: thicker/dark blue: exon, thinner/light blue: intron; coordinates relative to hg19.

We also found significant enrichment of these conditionally independent schizophrenia risk-associated eQTLs among genes with developmental isoform shits identified above – 44.0% of genes with eQTLs compared to 23.6% without eQTLs (OR=2.54, p=5.38x10^−8^). These conditional analyses could suggest potential regulatory roles of these unannotated transcribed sequences on annotated transcripts that play a putative role in the manifestation of schizophrenia risk in the brain. More generally, these eQTL results highlight significant and independently-replicated risk-associated schizophrenia candidate genes and their specific transcripts that comprise links in the causative chain of schizophrenia in the human brain.

### Expression associations with chronic schizophrenia illness

We lastly explored the expression landscape of the prefrontal cortex of the schizophrenia illness state and its potential link with developmental regulation and genetic risk. We performed differential expression modeling using 351 high quality adult samples (age >16, 196 controls, 155 cases), and found extensive bias by RNA degradation within both univariate analysis (where 12,686 genes were differentially expressed at FDR<5%) and even after adjusting for standard measured levels of RNA quality typical of all prior studies (Figure S6). We therefore implemented a novel statistical framework based on an independent molecular degradation experiment (see Methods, Results S3), called “quality surrogate variable analysis” (qSVA, see Methods)^18^. We further utilized potential replication RNA-seq data from the CommonMind Consortium (CMC) dataset, using a subset of age range-matched 159 schizophrenia patients and 172 controls. Interestingly, adjusting for observed factors related to RNA quality that characterize all earlier studies of gene expression in schizophrenic brain, including an earlier report using the CMC data ^16^, the proportion of genes with differentially expressed features at genome wide significant FDR < 5% that replicate (with directionality and marginal significance at p<0.05) in the CMC dataset was small (only 11.0%, 244/2,215). In contrast, using our new statistical qSVA approach, 40.1% of differentially expressed genes at FDR < 5% (N=75/183) replicate in the CMC dataset. At genome-wide significant FDR<10% (see Methods), we identified 237 genes with 556 DE features that replicated in the CMC dataset (33.6% gene-level replication rate, Table S9, Table S10).

The differences in expression levels between cases and controls of these DE features were generally small in both our discovery and the replication datasets (Figure 3A, Figure S8), perhaps a direct result of the clinical and molecular heterogeneity of this disorder ^13,19^. Gene ontology analysis implicated transporter- and channel-related signaling as significantly consistently downregulated in patients compared to controls across genes annotated in all three expression summarizations (Figure 3B, Table S11). These results suggested decreased signaling in patients with schizophrenia, but could raise the possibility that these replicated expression differences between patients and controls relate to epiphenomena of illness, such as treatment with antipsychotics which affect signaling in the brain^14^, as the majority of patients were on anti-psychotics at the time of death (64%, Table S1). Only two genes (*KLC1* and *PPP2R3A*) in the significant 108 schizophrenia GWAS loci were significantly differentially expressed. However, in an exploratory analysis, we found that overall the differential expression statistics within the loci were significantly different than those features outside the loci (Results S4, Figure S9, Table S12). We also investigated the relationships between transcription and genomic risk for schizophrenia using genome wide Polygene Risk Scores (PRS) from each subject calculated as previously described^2^ (see Methods). Using the subset of 209 Caucasian samples, we largely found a lack of association between PRS and expression of individual expression features. We further found a lack of enrichment of PRS on expression comparing the differentially expressed and replicated case-control features to the rest of the transcriptome, as well as lack of directionally consistency between PRS- and diagnosis-associated statistics among expressed features (Table S13). These results further suggest that the significant case-control expression differences show little overlap with genetic risk for the disorder.

**Figure 3:**
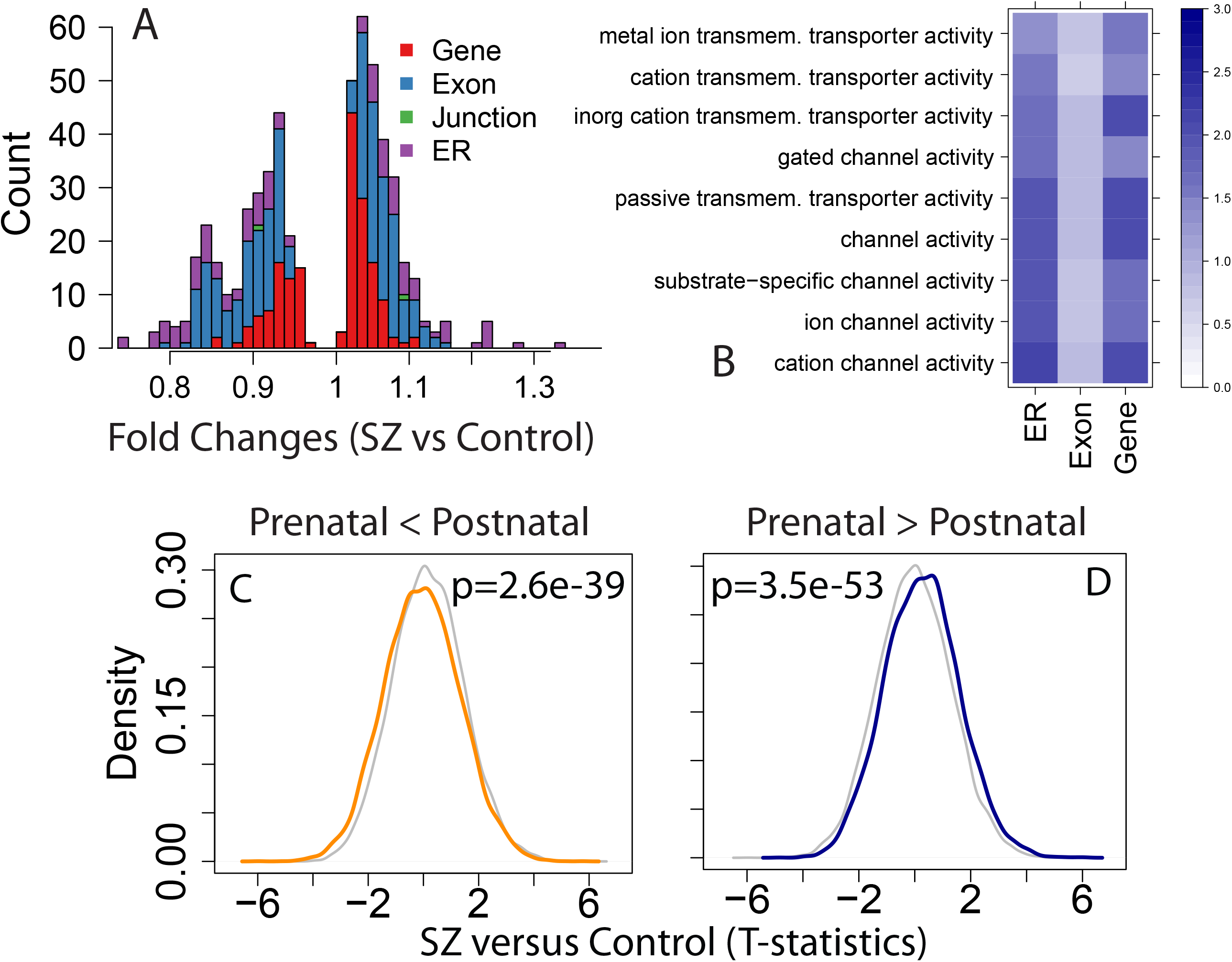
Differential expression comparing patients with schizophrenia to controls. (A) Histogram of fold changes of the diagnosis effect of those features that were significant and independently replicated, colored by feature type. (B) Gene set analyses of genes with decreased expression in patients compared to controls by feature type. Coloring/scaling represents -log10(FDR) for gene set enrichment. Significant directional effects of developmental regulation among diagnosis-associated features for those features that (C) increased and (D) decreased across development (i.e. those features shown in Figure 1B). P-values provided for Wilcoxon rank sign test for those features developmentally regulated among case-control differences to those not developmentally regulated.

In an earlier study of the epigenetic landscape of frontal cortex of patients with schizophrenia, we showed that DNA methylation levels in patients were closer to fetal methylation levels than to those of adult control samples^20^. Here we tested for analogous effects in the RNA-seq data related to the illness state. Every significant gene with differentially expressed features in the adult case-control analysis and replicated in the independent dataset showed evidence for developmental regulation across at least two expression feature types. We further found that expression features more highly expressed in postnatal life tended to be more lowly expressed in patients compared to controls (max: p=3.24x10^−11^, min: p=1.05x10^−70^, Figure 3C) and features more highly expressed in fetal life tended to be more highly expressed in patients with schizophrenia compared to controls (max: p=6.86x10^−33^, min: p < 10^−100^, Figure 3D). Analogous analyses for developmental regulation of schizophrenia-associated features without adjusting for the RNA quality qSVs were significant in the opposite directions, namely that schizophrenia-associated changes were further from, rather than closer to, fetal expression levels, as might be predicted as a confounding artifact of residual RNA quality differences (as the quality of the samples rank as fetal > adult control > adult SZ, see Table S1).These results further converge on a role for genes changing during brain development and maturation in the pathogenesis of schizophrenia, specifically that both DNA methylation and expression levels in adult patients appear to reflect levels in the developing brain more strongly than do those of unaffected individuals. These results also underscore the risk of spurious findings based on uncorrected RNA quality confounding.

## Discussion

We have explored the diverse landscape of expression correlates of schizophrenia risk and illness state in the postmortem human frontal cortex across the lifespan. Using deep RNA sequencing to define convergent measures of gene expression and early brain development, we identified widespread developmental regulation of transcription, including novel discoveries related to preferential isoform usage across brain development. These unexpected isoform “shifts” were associated with genetic risk for schizophrenia, and the directionality of dysregulation of developmentally regulated features suggest a more fetal-like expression profile in patients with schizophrenia compared with controls. Our approach to transcript characterization, which included extensive characterization of unannotated sequence, revealed that many more schizophrenia risk associated SNPs are brain eQTLs than previously reported - many risk SNPs only associate with a single gene, or even a single transcript, and many of these adult-identified eQTLs show overlap with genes with dynamic isoform regulation across human brain development. Lastly, we identified significant and replicated genes differentially expressed in patients with schizophrenia compared to unaffected controls using a new experiment-based statistical framework to estimate and reduce the effects of latent RNA degradation bias which had not been accounted for in earlier studies. Without this new approach to RNA quality adjustment, replication across datasets is markedly limited if not negligible, and the directionality of the association with developmental isoform shifts is anomalous. These data suggest a convergence of developmental regulation and genetic risk for schizophrenia that appears relatively stable in patients ascertained at death, following decades of illness after diagnosis. We previously observed analogous stability of epigenetic marks highlighting prenatal life in adult patients with schizophrenia^20^, suggesting that both genetic and environmental risk factors implicated in schizophrenia illness involve early developmental events that are still observable in the brain tissue of adult individuals despite many years of illness.

While our approach utilizing convergent expression features – genes, exons, transcripts, junctions, and expressed regions – results in more complicated data processing and analysis, it can potentially cast a wider net in the search for valid biological signals in RNA sequencing datasets. Using all convergent features overcomes the limitations related to any given feature summarization, including the inability to measure and interrogate unannotated or novel transcribed sequences using gene and exon counts, and the difficulties in full transcript assembly from short sequencing reads ^21^. We note that both quantifying and analyzing splice junctions, and also transcript-level data, rely on junction-spanning reads for statistical power. In our data, there were approximately 3 times (IQR: 2.86-3.24) more reads available by gene/exon counting approaches than those that contain splice junctions, likely explaining why gene counts discovered more differentially expressed genes in the schizophrenia diagnosis analyses. Two relatively new approaches utilized here – direct quantification and statistical analyses of splice junction counts and expressed regions – can identify differential expression signal when it is outside of the annotated transcriptome. The junction-level approach can also identify previously uncharacterized novel transcribed sequences, which we replicated in other large publicly available datasets, as well as delineate individual transcripts or classes of transcripts that share a particular splice junction. As read lengths increase, the proportion of reads containing splice junctions will increase, making junction- and transcript-based approaches even more powerful, including those recently developed to identify splicing QTLs ^22^.

Our analysis of RNA-seq data identified widespread shifts in preferential isoform use across brain development, which would have been impossible to identify using only gene-level data and incomplete with only exon-level data (Figure 2). The genes with these isoform shifts were significantly enriched for neurodevelopmental and cellular signaling processes, and as well as for genes in regions of genetic risk for schizophrenia. A prevalent hypothesis suggests that schizophrenia is a neurodevelopmental disorder that arises because of altered connectivity and plasticity in the early assembly of relevant neural circuits^23^, and the potential convergence of genetic risk with developing signaling processes across human brain development should point to specific candidate molecular disruptions occurring during the wiring of the fetal brain. Indeed, inefficient or disrupted signaling and tuning is thought to underlie the expression of illness in the adult brain ^23^, and the most successful therapeutics work through improving these processes^14^. Consistent with this hypothesis, we find evidence for differences in the expression of genes coding for subunits of ion channels in the cortices of patients with schizophrenia compared to controls. We observed significant differential expression of both voltage-gated (*KCNA1*, *KCNC3*, KCNK1, *KCNN1*, *SCN9A*) and ligand gated ion channels (*GRIN3A*, *GABRA5*, *GABRB3*), transporters (*SLC16A2*, *ALC25A33*, *SLC26A11*, *SLC35F2*, *SLC7A3*), and ion channel auxiliary subunits (*KCNIP3*, *SCN1B*), supporting other evidence that the clinical phenomenology of schizophrenia is associated with altered neuronal excitability ^24^. While these findings implicating basic mechanisms of cortical circuit dynamics may underlie fundamental aspects of the clinical disorder, the possibility that they are driven by the effects of pharmacological treatment and are thus state dependent epiphenomena cannot be excluded. Indeed, our failure to find association of genomic risk scores and differential gene expression in the illness state adds weight to the latter interpretation.

Our eQTL analyses are among the largest and most comprehensive to date in human brain tissue, based on stringent genome-wide significance and independent replication, and offer additional insights into the genetic regulation of RNA expression levels. Our data also suggest more widespread regulation of specific transcript isoforms, which we were able to identify using exon- and junction-level analyses. This transcript-specific genetic regulation was particularly prevalent among schizophrenia risk variants, where 66.9% of loci containing multiple transcripts showed clinically- and molecularly-consistent eQTL signal to a single Ensembl transcript isoform. Overall, we have identified many more eQTLs to genome-wide significant schizophrenia risk variants – 48.1% - than previously reported, experimentally implicating far more potential “risk” genes within these loci than previously characterized. Our database of eQTLs – available at eqtl.brainseq.org/phase1/eqtl – is searchable for candidate genes or SNPs and provides publication-ready visualizations (e.g. boxplots in Figure 2) and statistics eQTL associations. The database can serve as a “one stop shop” for eQTL statistics across three independent studies (LIBD, CMC, and GTEx) for both annotated and unannotated transcribed sequence in the human cortex, and can export results to the UCSC Genome Browser ^25^ for additional interrogation.

These eQTL associations within the genome-wide significant schizophrenia loci identify novel putative biological mechanisms underlying risk for the disorder. We have highlighted GWAS loci that contain significant and statistically independent eQTLs, as they often point to individual “risk” genes or even more specific “risk” transcripts. These “risk” genes and transcripts are targetable entry points for more focused cellular assays and model organism work to better characterize schizophrenia risk mechanisms. Moreover, these eQTLs of specific transcript features identifies a compelling strategy and directionality for target rescue, specifically to increase or decrease the function of the target transcript(s) and downstream effectors. Focusing solely on increased or decreased expression in brains of patients compared to controls, without considering genetic risk variants and their regulation of local gene expression, will likely predominantly highlight molecular changes resulting from the schizophrenia illness state, as we suggest with consistent down-regulation of ion channels. We stress the priority of identifying the most relevant cellular consequences of genetic risk, which we view as production of particular isoforms with predicted directionality, rather than trying to identify “causal” mutations tagged by “marker” risk SNPs from the GWAS. We suggest that identifying convergence between genetic risk and potential molecular consequences of the disorder is likely to result in better, or at least more consistent support for, targets for drug discovery efforts.

## Author Contributions

A.E.J – performed primary data processing and analyses, led the writing of the manuscript

R.E.S – contributed to data analysis and writing of the manuscript

J.H.S., R.T. , Y.G. – performed RNA sequencing data generation (RNA extraction, library preparation, and sequencing) and QC analyses

L.C.T.,J.T.L – performed region-level data generation and assisted in data analysis and interpretation

T.K.T.,S.X.,J.Q.,C.C., B.J.M., A.C.,N.B., BrainSeq – provided feedback on manuscript and contributed to data analyses and interpretations on eQTL analyses.

W.S.U. – created user-friendly database of eQTLs

A.D.S. – consented and clinically characterized human brain donors

T.M.H.,J.E.K.- collected, consented, characterized, and dissected human brain tissue; contributed to the design of the study

D.R.W. – designed and oversaw the research project, wrote the manuscript

Tony Kam-Thong is employed by F. Hoffmann-La Roche

Hualin S Xi and Jie Quan are employees of Pfizer Inc.

Alan Cross, and Nicholas J.Brandon were full time employees and shareholders in AstraZeneca at the time these studies were conducted.

The remaining authors declare no competing financial interests.

### Data Availability

sequencing reads and genotype data are available through SRA and dbGaP at accession numbers: [TBD] following publication.

## Acknowledgements

We thank Dr. Ronald Zielke, Robert D. Vigorito, and Robert M. Johnson of the National Institute of Child Health and Human Development Brain and Tissue Bank for Developmental Disorders at the University of Maryland for their provision of fetal, child, and adolescent brain specimens; This work was supported by the funding from Lieber Institute for Brain Development and the Maltz Research Laboratories. The work was partially supported by R21MH109956 to A.E.J. L.C.T was supported by Consejo Nacional de Ciencia y Tecnología México 351535.

The Genotype-Tissue Expression (GTEx) Project was supported by the <u>Common Fund</u> of the Office of the Director of the National Institutes of Health. Additional funds were provided by the NCI, NHGRI, NHLBI, NIDA, NIMH, and NINDS. Donors were enrolled at Biospecimen Source Sites funded by NCI\SAIC-Frederick, Inc. (SAIC-F) subcontracts to the National Disease Research Interchange (10XS170), Roswell Park Cancer Institute (10XS171), and Science Care, Inc. (X10S172). The Laboratory, Data Analysis, and Coordinating Center (LDACC) was funded through a contract (HHSN268201000029C) to The Broad Institute, Inc. Biorepository operations were funded through an SAIC-F subcontract to Van Andel Institute (10ST1035). Additional data repository and project management were provided by SAIC-F (HHSN261200800001E). The Brain Bank was supported by a supplements to University of Miami grants DA006227 & DA033684 and to contract N01MH000028. Statistical Methods development grants were made to the University of Geneva (MH090941 & MH101814), the University of Chicago (MH090951, MH090937, MH101820, MH101825), the University of North Carolina - Chapel Hill (MH090936 & MH101819), Harvard University (MH090948), Stanford University (MH101782), Washington University St Louis (MH101810), and the University of Pennsylvania (MH101822). The data used for the analyses described in this manuscript were obtained from <u>dbGaP</u> accession number <u>phs000424.v6.p1</u> on October 6, 2015.

Data were generated as part of the CommonMind Consortium supported by funding from Takeda Pharmaceuticals Company Limited, F. Hoffman-La Roche Ltd and NIH grants R01MH085542, R01MH093725, P50MH066392, P50MH080405, R01MH097276, RO1-MH-075916, P50M096891, P50MH084053S1, R37MH057881 and R37MH057881S1, HHSN271201300031C, AG02219, AG05138 and MH06692. Brain tissue for the study was obtained from the following brain bank collections: the Mount Sinai NIH Brain and Tissue Repository, the University of Pennsylvania Alzheimer’s Disease Core Center, the University of Pittsburgh NeuroBioBank and Brain and Tissue Repositories and the NIMH Human Brain Collection Core. CMC Leadership: Pamela Sklar, Joseph Buxbaum (Icahn School of Medicine at Mount Sinai), Bernie Devlin, David Lewis (University of Pittsburgh), Raquel Gur, Chang-Gyu Hahn (University of Pennsylvania), Keisuke Hirai, Hiroyoshi Toyoshiba (Takeda Pharmaceuticals Company Limited), Enrico Domenici, Laurent Essioux (F. Hoffman-La Roche Ltd), Lara Mangravite, Mette Peters (Sage Bionetworks), Thomas Lehner, Barbara Lipska (NIMH).

Members of the BrainSeq consortium include: Christian R. Schubert, Patricio O’Donnell, Jie Quan, Jens R. Wendland, Hualin S. Xi, Ashley R. Winslow, Enrico Domenici, Laurent Essioux, Tony Kam-Thong, David C. Airey, John N. Calley, David A. Collier, Hong Wang, Brian Eastwood, Philip Ebert, Yushi Liu, Laura Nisenbaum, Cara Ruble, James E. Scherschel, Ryan Matthew Smith, Hui-Rong Qian, Kalpana Merchant, Michael Didriksen, Mitsuyuki Matsumoto, Takeshi Saito, Nicholas J. Brandon, Alan J. Cross, Qi Wang, Husseini Manji, Hartmuth Kolb, Maura Furey, Wayne C. Drevets, Joo Heon Shin, Andrew E. Jaffe, Yankai Jia, Richard E. Straub, Amy Deep-Soboslay, Thomas M. Hyde, Joel E. Kleinman, Daniel R. Weinberger

## Methods

### Postmortem brain samples

Post-mortem human brain tissue was obtained by autopsy primarily from the Offices of the Chief Medical Examiner of the District of Columbia, and of the Commonwealth of Virginia, Northern District, all with informed consent from the legal next of kin (protocol 90-M-0142 approved by the NIMH/NIH Institutional Review Board). Additional post-mortem fetal, infant, child and adolescent brain tissue samples were provided by the National Institute of Child Health and Human Development Brain and Tissue Bank for Developmental Disorders (http://www.BTBank.org) under contracts NO1-HD-4-3368 and NO1-HD-4-3383. The Institutional Review Board of the University of Maryland at Baltimore and the State of Maryland approved the protocol, and the tissue was donated to the Lieber Institute for Brain Development under the terms of a Material Transfer Agreement. Clinical characterization, diagnoses, and macro- and microscopic neuropathological examinations were performed on all samples using a standardized paradigm, and subjects with evidence of macro- or microscopic neuropathology were excluded. Details of tissue acquisition, handling, processing, dissection, clinical characterization, diagnoses, neuropathological examinations, RNA extraction and quality control measures were described previously in Lipska, et al. ^26^. The Brain and Tissue Bank cases were handled in a similar fashion (http://medschool.umaryland.edu/BTBank/ProtocolMethods.html). Antipsychotic use was measured using toxicology at time of death.

### RNA extraction and sequencing

Post-mortem tissue homogenates of dorsolateral prefrontal cortex grey matter (DLPFC) approximating BA46/9 in postnatal samples and the corresponding region of PFC in fetal samples were obtained from all subjects. Total RNA was extracted from ~100 mg of tissue using the RNeasy kit (Qiagen) according to the manufacturer’s protocol. The poly-A containing RNA molecules were purified from 1 µg DNAse treated total RNA and sequencing libraries were constructed using the Illumina TruSeq© RNA Sample Preparation v2 kit. Sequencing indices/barcodes were inserted into Illumina adapters allowing samples to be multiplexed in across lanes in each flow cell. These products were then purified and enriched with PCR to create the final cDNA library for high throughput sequencing using an Illumina HiSeq 2000 with paired end 2x100bp reads.

### RNA sequencing data processing

The Illumina Real Time Analysis (RTA) module performed image analysis, base calling, and the BCL Converter (CASAVA v1.8.2), generating FASTQ files containing the sequencing reads. These reads were aligned to the human genome (UCSC hg19 build) using the spliced-read mapper TopHat (v2.0.4) using the reference transcriptome to initially guide alignment, based on known transcripts of the previous Ensembl build GRCh37.67 (the “–G” argument in the software) ^27^. We achieved a median of 85.3 million (IQR: 71.7M-111.2M) aligned reads per sample (see Table S1).

We characterized the transcriptomes of these 495 samples using five convergent measurements of expression (“feature summarizations”)– (1) gene and (2) exon counts, and (3) transcript-level quantifications that rely on existing gene annotation, and two annotation-agnostic approaches we have developed that are determined solely from the read alignments – (4) read coverage supporting exon-exon splice junctions (e.g. coordinates of potentially intronic sequence that are spliced out of mature transcripts captured by a single read) and (5) read coverage overlapping each base in each sample which we have summarized into contiguous “expressed regions” (ERs, see Methods, Figure S1). These last three measurements generate expression for features of interest that can “tag” elements of transcripts in the data that are not constrained by limitations or incompleteness of existing annotation, and the counts for these features can then be directly used for differential expression analysis.

1. Gene counts were generated using the featureCounts tool^28^ (v1.4.3-p1) based on the more recent Ensembl v75, which was the last stable release for the hg19 genome build, using single end read counting [featureCounts –a $GTF –o $OUT $BAM]. We converted counts to RPKM values using the total number of aligned reads across the autosomal and sex chromosomes (dropping reads mapping to the mitochondria chromosome).
2. Exon counts were also generated using the featureCounts tool^28^ (v1.4.3-p1) based on the more recent Ensembl v75, using single end read counting, and allowing reads to be assigned to multiple exons (e.g. those with splice junctions) [featureCounts –O –f –a $GTF – o $OUT $BAM]. We converted counts to RPKM values using the total number of aligned reads across the autosomal and sex chromosomes (dropping reads mapping to the mitochondria chromosome).
3. Junction counts were generated by first filtering the TopHat BAM file to primary alignments only [samtools view -bh -F 0x100 $BAM > $NEWBAM ] and regtools ^29^ (v 0.1.0) was used to extract analogous junction information (coordinates and number of reads supporting) as the TopHat output. We found that native TopHat output (junctions.bed) was based on both primary and secondary alignments, which could influence the degree of potentially novel splice junctions. We used a modified version of TopHat’s “bed_to_juncs” program to retain the number of supporting reads (in addition to returning the coordinates of the spliced sequence, rather than the maximum fragment range), and used R code (see Supplementary Code) to combine and annotate these junctions across all samples. We identified splice junctions using Ensembl v75 – while the initial alignment was guided by Ensembl v67, novel junctions, by definition, are identified in the second genome alignment, rather than the initial guided transcriptome alignment step. We converted counts to “RP80M” values, or “reads per 80 million mapped” using the total number of aligned reads across the autosomal and sex chromosomes (dropping reads mapping to the mitochondria chromosome), which can be interpreted as the number of reads supporting the junction in an average library size (we were targeting 80M reads in the sequencing). Most junctions were lowly expressed in our homogenate tissue, with fewer than 1 average normalized supporting read (N=3,330,642; 92.98%) including approximately half unique to a single individual (N= 1,779,241, 49.67%).
4. Transcripts were assembled using StringTie^4^ (version 1.1.2) guided by Ensembl v75 annotation within each sample [stringtie $BAM –o $OUT –G $GTF]. We then used “CuffMerge” ^30^ to merge all assembled transcriptomes across all samples, and then re-quantified the expression of each transcript isoform in each sample again using StringTie to this global set of transcripts [stringtie $BAM –B –e –o $OUT –G $GTF_ALL] to have expression measurements on the same transcripts across all samples. We then used the “ballgown” tool^31^ to merge all assembled and quantified transcripts across all samples (N=733,339), and used liberal filtering to remove lowly or uniquely expressed transcripts (mean FPKM > 0.025), resulting in 188,578 transcripts across the 495 samples.
5. Expressed regions (ERs) were calculated using the “derfinder” R Bioconductor package^6^ using a cutoff of 5 normalized (to 80M reads) read coverage, which identified 389,797 ERs. We retained the 275,885 ERs that were at least 12 basepairs, and annotated the ERs to Ensembl v75.

### Genotype data processing

SNP genotyping with HumanHap650Y_V3 (N=135), Human 1M-Duo_V3 (N=357), and Omni5 (N=3) BeadChips (Illumina, San Diego, CA) was carried out according to the manufacturer’s instructions with DNA extracted from cerebellar tissue. Genotype data were processed and normalized with the crlmm R/Bioconductor package^32^ separately by platform. Genotype imputation was performed on high-quality observed genotypes (removing low quality and rare variants) using the prephasing/imputation stepwise approach implemented in IMPUTE2^33^ and Shape-IT^34^, with the imputation reference set from the full 1000 Human Genomes Project Phase 3 data set, separately by platform. We retained common SNPs (MAF > 5%) that were present in the majority of samples (missingness < 10%) that were in Hardy Weinberg equilibrium (at p > 1x10^−6^) using the Plink^35^ version 1.9 tool kit [‘plink --bfile $BFILE --geno 0.1 --maf 0.05 --hwe 0.000001’]. We then identified linkage disequilibrium (LD)-independent SNPs to use in genome-wide clustering of samples and in the number of independent eQTL tests performed [‘plink – bfile $BFILE --indep 100 10 1.25’]. Multidimensional scaling (MDS) was performed on the autosomal LD-independent construct genomic ancestry components on each sample, which can be interpreted as quantitative levels of ethnicity – the first component separated the Caucasian and African American samples. This processing and quality control steps resulted in 7,421,423 common variants in this dataset of 495 subjects.

#### Polygene risk score (PRS) analysis

Using the allelic dosage files following imputation described above and the SNPs from provided by the PGC to the Lieber Institute that did not contain completely different clinical subjects used in the GWAS^2^. We considered expression associations at the gene, exon and junction-level to the PRS scores from the first 5 clinical SNP sets, corresponding to GWAS p-value thresholds of p < 5e-8 (s1), p < 1e-6 (s2), p < 1e-4 (s3), p < 0.001 (s4), and p < 0.01 (s5) – subsequent SNP sets were ignored due to clinical risk plateauing at s5. We also focused only on Caucasian individuals (96 cases, 113 controls), as the s5 PRS was increased in patients relative to controls in this sample (p=3.2x10^−5^), but did not differ among African Americans (p=0.9). Within each expression feature type, we modeled expression levels as a function of each PRS set (s1-s5), adjusting for 3 MDS components of the genotype data, sex, and the first *K* principal components (PCs) of the normalized expression features, where *K* was calculated using the Buja and Eyuboglu permutation-based algorithm^36^ in the “sva” Bioconductor package^37^. The resulting p-values of PRS on expression, adjusting for the above factors, were subject to false discovery rate (FDR) control to account for multiple testing.

### Public data processing

#### GTEx

Raw RNA-seq reads from all brain samples with corresponding genotype data were downloaded from SRA and aligned to the genome using TopHat2 ^27^ (version 2.0.14) using the iGenomes transcriptome and genome annotations based on hg19. As above, featureCounts ^28^ was used to quantify expression of genes and exons relative to Ensembl v75, and junctions were quantified with regtools^29^ as above. We used StringTie with the assembled merged GTF from the LIBD DLPFC samples on the GTEx BAM files to quantify the same transcripts, and used bwtool^38^ to quantify the coverage of the same expressed regions from the GTEx brain samples. Genotype data from the two platforms (Illumina Omni 5M and 2.5M) were imputed separately as described above and merged into a single plink^35^ set.

#### GEUVADIS

Raw RNA-seq reads from all LCL samples were downloaded from SRA and aligned to the genome using TopHat2 ^27^ (version 2.0.9) using the iGenomes transcriptome and genome annotations based on hg19. As above, featureCounts^28^ was used to quantify expression of genes and exons relative to Ensembl v75, and junctions were quantified with regtools^29^ as above. We used StringTie with the assembled merged GTF from the LIBD DLPFC samples on the GEUVADIS BAM files to quantify the same transcripts, and used bwtool to quantify the coverage of the same expressed regions from the GEUVADIS LCL samples.

#### CommonMind Consortium (CMC)

547 BAM files were downloaded from Synapse, which were aligned with TopHat2 (version 2.0.9) using Ensembl v70 transcriptome annotation and the hg19 genome. As above, featureCounts ^28^ was used to quantify expression of genes and exons relative to Ensembl v75, and junctions were quantified with regtools ^29^ as above. We used StringTie with the assembled merged GTF from the LIBD DLPFC samples on the CMC BAM files to quantify the same transcripts, and used bwtool to quantify the coverage of the same expressed regions from the CMC brain samples. Genotypes were converted to plink file sets from GEN files obtained from Synapse using posterior probabilities > 90%, resulting in genotype data across 9,506,038 SNPs and 547 samples.

### Differential expression across brain development

We modeled differential expression across age at each of the five feature summarizations (gene, exon, junction, transcript, and ER) in the 320 control subjects across the lifespan. We modeled expression, after transforming with log2 with an offset of 1, as a function of age after creating using linear splines with breakpoints at ages: birth (0), 1, 10, 20, and 50, further adjusting for sex and ancestry/ethnicity (first 3 MDS components). F-statistics were computed comparing the model containing age (including the linear splines), sex, and ethnicity, to a statistical model with just sex and ethnicity, with corresponding p-values calculated based on an F-distribution with 11 and 308 degrees of freedom, and Bonferroni adjustment within each feature type was performed using the number of features with non-zero expression (gene RPKM > 0.01, exon RPKM > 0.1, and junction RP80M > 0.2 with non-novel annotation) across all samples as the number of tests (which varied by feature type). We also computed post-hoc statistics on the data, including the Pearson correlation between “cleaned” expression (after regressing out the effects of sex and ethnicity, holding the age effects constant), and age to determine if the expression of the fetal rose or fell across the lifespan, and also measured the fetal versus postnatal log_2_ fold changes.

Preferential isoform usage across aging was determined by identifying the subset of genes (by Ensembl ID) that contained at least one Bonferroni-significant feature that had positive correlation with age and another Bonferroni-significant feature that had negative correlation with age. We also computed the difference in positive and negative correlations as a measure of the magnitude of the preferential isoform use. Gene set analyses using pre-defined gene ontology (GO) and Kyoto Encyclopedia of Genes and Genomes (KEGG) sets were performed using the clusterProfiler R/Bioconductor package^39^, here using the genes (mapping from Ensembl to Entrez ID) that had such preferential isoform use to those that were developmentally regulated (having at least one feature that was associate with age at Bonferroni significance). Enrichments with the PGC2 schizophrenia risk loci – defined by the chr:start-end roughly corresponding to linkage disequilibrium blocks in the published manuscript - were performed both parametrically, by overlapping the genomic coordinates of the 108 risk regions with those genes that had preferential isoform usage, compared to a background of all genes with each set of expressed features, as well as by permuting the locations of the 108 regions across the genome 10,000 times and each time, re-computing the overlap within these null regions – see additional details in Jaffe et al 2015^7^. Empirical p-values were calculated by counting the number of the odds ratios across the 10,000 null permutations to each observed odds ratio.

### eQTL discovery analyses

We performed eQTL analyses separately by feature type (gene, exon, junction, transcript, and ER) allowing for a 500kb window around each of the 7,421,423 common SNPs in the 412 age > 13 samples, adjusting for ancestry (first three MDS components from the genotype data), sex, diagnosis, and the first *K* principal components (PCs) of the normalized expression features, where *K* was calculated separately by feature type using the Buja and Eyuboglu permutation-based algorithm^36^ in the “sva” Bioconductor package^37^ (gene: 22 PCs, exon: 19 PCs, junction: 26 PCs, transcript: 25 PCs, expressed regions: 20 PCs). The eQTL analyses were run using the MatrixEQTL R package^40^, which returned the log_2_ fold change per allele copy, and corresponding T-statistic, p-value, and FDR for each SNP-feature pair. We further used the LD-independent SNPs to estimate the effective number of tests (by counting the number of features within a 500kb window around each LD independent SNP) for a more conservative Bonferroni adjustment. For all five feature types, we retained all eQTLs with FDR < 1%.

### eQTL replication analyses

We sought to replicate all significant SNP-Feature pairs for each eQTL in two independent datasets across all five feature summarizations: CommonMind Consortium and the GTEx project. We used chromosome and position of variants to attempt to match across dataset – almost all SNPs in the discovery sample were present in each replication samples. Within each dataset, we tested all polymorphic SNPs (e.g. not monomorphic) and corresponding expressed features, adjusting for the first 10 PCs of each feature summarization type and the first 5 MDS components of the corresponding common genotype data. Analyses within CMC were performed on the 285 controls and analyses in GTEx were performed within each brain region separately. After identifying and matching back on SNP-feature eQTL pairs, we checked whether the counted alleles were the same within the discovery and replication datasets and flipped the directionality of eQTL associations where the alleles were discordant. Note that in GTEx, some residual discordancy was still present across dataset (e.g. off-diagonal points in Figures S6, S7A and S7B) but not within a dataset (Figure S7C). Meta-analysis between discovery (LIBD) and CMC was performed using Stouffer’s Methods ^41^, by summing the T-statistics and dividing by the square-root of the number of datasets (N=2). Meta-analysis within GTEx brain regions was performed using the same approach, here dividing by the square root of number of datasets/brain regions (N=13). When replication statistics were not present in replication datasets due no/low expression or being monomorphic, the discovery eQTL was “penalized” by setting the replication statistic to 0 prior to meta-analysis.

### eQTL clinical enrichment analyses

We downloaded the 128 linkage-disequilibrium-independent variants that reached genome-wide significance in combined analysis from the latest schizophrenia GWAS (their Supplementary Table 2) and matched those variants to our data by chromosome and position relative to hg19. Of the 128 variants, only 106 were present in our final QC’d and common (MAF>5%) genotype data. Most were excluded due to MAFs less than 5% although several variants were dropped for other reasons (not present in 1000 Genomes, failed Hardy Weinberg equilibrium, poorly imputed, etc). We therefore interrogated only those 106 schizophrenia-associated variants among our eQTL associations. We utilized a similar strategy for the latest NHGRI GWAS catalog (downloaded 7/24/2017) with an additional step of lifting over our variants to hg38 and again matching by variant coordinates. Here, only approximately half of the variants were well-measured in our samples (see Table 2).

### eQTL conditional analyses

We performed conditional analyses within the eQTLs for each schizophrenia risk variant to remove highly correlated signal and improve resolution of associations. We used the residuals of the statistical model described above within each feature type (regressing out PCs, MDS components and diagnosis) to allow for analyses across feature types. We iteratively conditioned on the expression level of the most significant eQTL feature and recomputed the eQTL p-values for all other features to the risk SNP. Those features that were still marginally significantly (at p<0.05) were retained, and then next-best expression feature (following conditioning) was additionally adjusted for in the statistical model. This procedure of iteratively testing for conditional independence among remaining features and subsequently adjusting for the most significant feature continued until no additional features were independently associated with the genetic risk variant at p < 0.05. This procedure was performed separately within each of the 51 loci with eQTL signal.

### Schizophrenia differential expression analyses

#### Discovery dataset analysis

we first filtered the subjects with RNA-seq to retain a more stringent set of 155 SCZD cases and 196 controls (criteria: ages between 17-80, gene assignment rate > 0.5, mapping rate > 0.7, RIN > 6, not outlying on 2nd ancestry PC, only self-reported Caucasians and African Americans). We fit three statistical models across each of the expression summarizations, modeling log_2_ transformed expression (with an offset of 1) as a function of:

1. Adjusted (“_adj“ suffix in supplementary tables): SCZD diagnosis, adjusting for age, sex, ancestry (SNP PCs 1, 5, 6, 9, 10, which were at least marginally associated with diagnosis), and then observed measures related to RNA quality: RIN, mitochondrial mapping rate, and gene assignment rate.
2. Adjusted + Quality Surrogate Variables (“_qsva” suffix in supplementary tables): SCZD diagnosis adjusting for “Adjusted” model as well as the first 12 PCs from the degradation matrix (see below) based on polyA+ libraries (selected using to using the BE algorithm ^36^ in the sva Bioconductor package^37^ while providing the adjusted model as input).
3. Adjusted + Principal Components (“_pca” suffix in supplementary tables): SCZD diagnosis adjusting for "Adjusted" model as well as the first *k* PCs from the expressed features (using the 50000 most variable features) depending on the feature type (gene: 23 PCs, exon: 20 PCs, transcript: 26 PCs, junction: 26 PCs, ERs: 23 PCs).

We used the ‘lmTest’ and ‘ebayes’ functions in the limma Bioconductor package ^42^ to fit all ofthe statistical models to estimate log_2_ fold changes, moderated T-statistics, and correspondingp-values. Multiple testing correction via the false discovery rate (FDR) was applied using the set of expressed features in this sample set for each summarization type: 24,122 genes (mean RPKM > 0.1), 420,022 exons (mean RPKM > 0.2), 61,950 transcripts (mean FPKM > 0.2), 229,846 junctions (mean RP80M > 1), and the 275,885 ERs.

#### RNA quality correction

We summarize the RNA quality correction approach here – for more detail, see the companion paper by Jaffe et al 2017. Briefly, the quality surrogate variable analysis (qSVA) uses RNA sequencing data generated from five DLPFC tissue samples left unfrozen for 0, 15, 30 and 60 minutes, resulting in 20 RNA samples. These samples were sequenced with both polyA+ and RiboZero library preparations, and gene, exon and junction counts were derived as above. We utilized the gene-level effects of degradation in these data in Figure S5 to demonstrate residual confounding by RNA quality, which we call the “DEQual Plot”.

For a given preparation type, we identified the genomic regions most susceptible to degradation by correlating coverage at expressed regions ^6^ to degradation time, adjusting for donor. This statistical modeling identified 515 regions significantly susceptible to degradation (at Bonferroni significance) in the RiboZero libraries and the top 1000 regions most susceptible to degradation (among the 35,287 at Bonferroni significance) in the polyA+ libraries – the BED files for these degradation-susceptible regions are available in Jaffe et al 2017^18^

The algorithm then involves selecting the set of regions for a particular library type and calculating total coverage within each region in the new user-provided samples (e.g. the 495 DLPFC RNA-seq polyA+ samples) to form the degradation matrix (which is either 515 or 1000 rows by N samples). Then PCA is performed on the log2 transformed degradation matrix (with an offset of 1) and the top *K* PCs are selected, for example using the BE algorithm ^36^, and extracted – the set of these PCs are referred to as quality surrogate variables (qSVs), and are included as adjustment variables in subsequent differential expression analyses.

#### Replication dataset analysis

we performed analogous sample selection procedures as in the discovery dataset to select 159 patients and 172 controls (total gene assignment rate > 0.3, alignment rate > 0.8, RIN > 6, ages between 18-80, non-outlying on genetic ancestry PCs 3 and 5 and keeping only reported Caucasians and African Americans). We similarly fit the three sets of statistical models to all five feature summarizations, with the following differences compared to the discovery analysis:

1. Adjusted model: the model here was diagnosis adjusting for age, sex, race, brain bank, RIN, gene assignment rate, alignment rate.
2. qSVA model: the degradation matrix was constructed using the 515 regions based on the RiboZero libraries in the degradation experiment.
3. PC adjustment: for each feature summarization type, we included: 27 PCs for genes, 29 PCs for exons, 39 PCs for transcripts, 39 PCs for junctions, and 33 PCs for ERs.

In these replication data we did not perform FDR correction. We were using the study for replication, not discovery, and therefore only used the features that were expressed in our data regardless of the expression levels in CMC. We considered features independently replicated if they had the same directionality for the SCZD versus control log_2_ fold change and were marginally significant (at p < 0.05) in the CMC dataset.

Gene set analyses on replicated differentially expressed features and genes were performed with clusterProfiler^39^ as described above. Set-level analyses on features in the GWAS risk regions were conducted by assigning each expressed feature a binary variable for whether it was in the risk regions or not. Then we fit a linear regression model of the t-statistics for diagnosis, adjusted by the qSVA approach, as a function as whether the feature was in the risk region, adjusting for its average expression level. This analysis was conducted across and then within each of the five feature summarization types.

## Author information

sequencing reads and genotype data are available through SRA and dbGaP at accession numbers: [**TBD**].

Correspondence and requests for materials should be addressed to Andrew Jaffe. The following authors have competing interests: Tony Kam-Thong is employed by F. Hoffmann-La Roche Hualin S Xi and Jie Quan are employees of Pfizer Inc.

The remaining authors declare no competing financial interests.

## Supplementary Information

### Supplementary Figure Legends

**Figure S1**: Study overview and cartoon describing quantifying the five different expression summarizations.

**Figure S2**: Cartoon describing the four different splice junction annotation classes, relative to annotated exons (dark blue rectangles). (A) Annotated splice junctions map between two exons in a known transcript. (B) Exon-skipping splice junctions map to two annotated exons in different transcripts. (C) Alternative start/exon junctions map to only one annotated exon on either the 5’ or 3’ end. (D) Completely novel junction do not map to any known exon.

**Figure S3**: Venn diagram of developmentally regulated features mapped back to Ensembl Gene IDs by the five feature summarization methods. DER: differentially expressed region.

**Figure S4:** Example of *CRTC2* (A) containing a developmental isoform shift. (B) Gene-level analysis shows no developmental regulation but at the junction-level (C) one splice junction significantly decreases in expression and (D) another splice junction significantly increases in expression over the lifespan. Exons in panels (E), (F), and (H) show some marginal increases in expression across the lifespan, but only the exon in (G) is unique to a single isoform and shows significant decreases in expression.

**Figure S5**: Venn diagram of Ensembl Gene IDs that contain significant isoform shifts by the four feature summarization methods that allow for multiple features per gene. DER: differentially expressed region.

**Figure S6**: Discovery (LIBD) and replication (CMC) T-statistics for eQTLs identified in the DLPFC for the best SNP-feature pair for each feature across 5 feature summarization types.

**Figure S7**: Assessing regional specificity of eQTLs in GTEx for the best SNP-feature pair for each feature across 5 feature summarization types. (A) Significant replication of many eQTLs within discovery (LIBD) and Frontal Cortex samples. (B) These DLPFC-identified eQTLs showed very significant meta-analysis T-statistics across the 13 brain regions in GTEx. (C) These DLPFC-identified eQTLs showed lack of regional specificity even within GTEx.

**Figure S8**: Scatter plot of effect sizes (fold changes) in discovery and replication datasets for those features significant and replicated. Colors have the same legend as Figure 3A.

**Figure S9**: GWAS loci set-level analysis for (A) all features together and then stratified by only (B) genes, (C) exons, (D) junctions, (E) transcripts and (F) expressed regions. P-values were based on the Wilcoxon rank sign test.

### Supplementary Table Legends

**Table S1**: Demographic information for subjects in the present study, stratified by age and diagnosis group. Dx: diagnosis, N: sample size, F: Female, Cauc: Caucasian, SD: standard deviation, PCW: post-conception weeks. Antipsychotic use was measured using toxicology at time of death. P-values for diagnosis differences in continuous variables are based on linear regression and P-values for categorical variables are based on chi-squared tests.

**Table S2**: Splice junction annotation and characterization in GTEx and GEUVADIS for any junction or highly expressed junctions (mean reads per 80M mapped reads, RP80M > 0, > 1 and > 5). Each column represents a 2x2 table for presence of identified junctions in 495 DLPFC samples in two independent polyA+ datasets.

**Table S3**: Summary statistics for those features significantly developmentally regulated in the control-only analyses across the lifespan.

**Table S4**: Significant developmentally regulated features collapsed to Ensembl Gene ID, used to make Figure S3

**Table S5**: Isoform shifts by Ensembl Gene ID and feature summarization type.

**Table S6**: Gene set analyses for those genes with significant isoform shift, stratified by feature summarization type. Q-values, which control the false discovery rate, FDR, are shown.

**Table S7**: Genes within the PGC schizophrenia GWAS risk regions that contain isoform shifts by feature summarization type. 21.8% of PGC2 genes had developmental isoform shifts using exon counts (N=96/440) and 31.9% showed this isoform shift association based on junction counts (N=137/430)

**Table S8**: Significant eQTLs to schizophrenia GWAS index variants, including replication statistics and additional annotation metrics for variants and expressed features. “condIndep” column refers to those associations that were conditionally independent.

**Table S9**: Differential expression statistics for those features that were significant and replicated in case-control comparisons.

**Table S10**: Genes consistently differentially expressed by case-control analysis for the different feature summarizations.

**Table S11**: Gene set analysis for genes with features differentially expressed by case-control status, stratified by directionality and feature summarization type.

**Table S12**: GWAS region set-level analyses for diagnosis-associated differentially expressed features, testing whether features in the PGC risk loci were more or less expressed as a set in cases compared to controls. Qual: qSVA adjusted analysis, Adj: observed covariate adjusted analysis.

**Table S13**: Associations between diagnosis, RPS and expression at gene and exon levels. First two columns for each feature: p-values for gene set tests for the significant case-control features among statistics capturing the effect of RPS on expression. Second two columns for each feature: directionality between RPS on expression associations and diagnosis on

